# Reduced entropy of subthalamic beta bursts predicts freezing of gait in Parkinson’s disease

**DOI:** 10.64898/2026.08.13.744293

**Authors:** Christopher A. Beaudoin, Andrew B. O’Keeffe, Bahman Abdi-Sargezeh, Martin J. Gillies, Ashwini Oswal, Alexander L. Green

## Abstract

**Background:** Freezing of gait (FOG) in Parkinson’s disease is associated with abnormal beta activity in the subthalamic nucleus (STN), but the temporal structure of burst dynamics remains poorly understood.

**Objectives:** To determine whether temporal features of STN beta bursts distinguish pre-freeze from stable gait and predict freezing onset.

**Methods:** STN recordings during gait from four individuals were analyzed. Temporal features of burst timing, including entropy and variability, were computed across behavioral states. Predictive performance was assessed using leave-one-patient-out classifiers.

**Results:** Entropy of inter-burst intervals was reduced prior to freezing (p < 0.01), with strong predictive performance (AUC = 0.825; threshold AUC = 0.858). During freezing, variability measures decreased and temporal structure increased, while entropy did not differ from pre-freeze. Phase-amplitude coupling showed frequency-specific but heterogeneous effects across comparisons.

**Conclusions:** Reduced temporal variability of STN beta burst timing precedes and predicts freezing, suggesting a transition to constrained neural dynamics.

## Introduction

Freezing of gait (FOG) is a disabling symptom of Parkinson’s disease characterized by transient episodes in which patients are unable to initiate or continue stepping(1). These events often occur during turning, increasing fall risk and reducing quality of life(2). Increasing evidence suggests that FOG reflects impaired coordination across distributed locomotor networks(3–5). Abnormal beta and theta oscillatory activity in the subthalamic nucleus (STN), including prolonged beta bursts, has been linked to FOG(6–11). Burst-based measures appear to capture motor impairment more effectively than averaged beta power(12). However, most studies have focused on burst duration or rate, leaving the broader temporal organization of burst timing largely unexplored(13). Although entropy, a measure of the unpredictability of a signal, differs between individuals with and without FOG(14), it remains unclear whether such changes emerge within individuals prior to freezing events.

Here, temporal dynamics of STN beta bursts were analyzed during gait. Entropy-based measures were used to quantify temporal complexity, alongside variability and cross-frequency coupling metrics. Phase-amplitude coupling (PAC), a measure of cross-frequency coordination, has been implicated in FOG, with abnormal coupling patterns observed in cortical recordings and modulated by deep brain stimulation(15). Analyses were performed on STN recordings obtained by Thenaisie *et al*.*(16)* during gait in four individuals with Parkinson’s disease, enabling detailed within-subject comparisons across behavioral states, including FOG. The study addressed three questions: whether burst dynamics distinguish pre-freeze from stable turning, how these dynamics change during freezing, and whether these features can predict freezing onset.

## Methods

STN local field potential recordings during gait were analyzed from four individuals with Parkinson’s disease from a previously published dataset(16) (clinical details in Table S1). For each session, a single STN contact was selected as the channel with the highest beta power, following the approach of Thenaisie *et al*.*(16)*. Beta bursts were defined from the Hilbert envelope of band-pass filtered signals using a 75th percentile threshold and minimum duration criterion and assigned to behavioral states based on burst midpoint timing. For each turn-freeze cluster, the turn preceding the first freeze bout was defined as the pre-freeze turn, while turns not adjacent to freezing were classified as stable (Table S2). Temporal features of burst timing were computed within each behavioral bout, including variability and complexity measures derived from inter-burst intervals, and Shannon entropy of burst timing computed from the distribution of burst midpoint times. Predictive performance was assessed using leave-one-patient-out logistic regression and threshold classifiers. Full methodological details are provided in the Supplementary Material.

## Results

### Reduced entropy precedes freezing and predicts freezing onset

Most freezing episodes were preceded by turning (163/176, 92.6%), approaching 100% in three of four patients. Accordingly, burst timing during turning episodes preceding freezing (pre-freeze turns) was compared with turning not associated with freezing (stable turns) at the level of individual turning episodes (clusters). Shannon entropy of burst timing was reduced during pre-freeze turns in both low and high beta bands (high beta: p = 0.00186, low beta: p = 0.000567; Figure 1B). Entropy was lower in pre-freeze than stable turns (Δ = 0.611 ± 0.282). This effect remained significant after adjusting for bout duration in a linear mixed-effects model (β = -0.529, p = 0.0027), while bout duration was not associated with entropy (β = 0.047, p = 0.561). Lempel-Ziv complexity, computed after binarization of inter-burst intervals, was increased (0.222 ± 0.087) during pre-freeze turns (high beta: p = 0.000621; low beta: p = 0.00723), consistent with increased temporal structure (Figure 1C). No interaction was observed between condition and beta sub-band for entropy (p = 0.693) or Lempel-Ziv complexity (p = 0.725). No statistical significance was found for burst rate, time-in-burst, coefficient of variation (CV), median absolute deviation (MAD), sample entropy, or permutation entropy.

**Figure 1.**
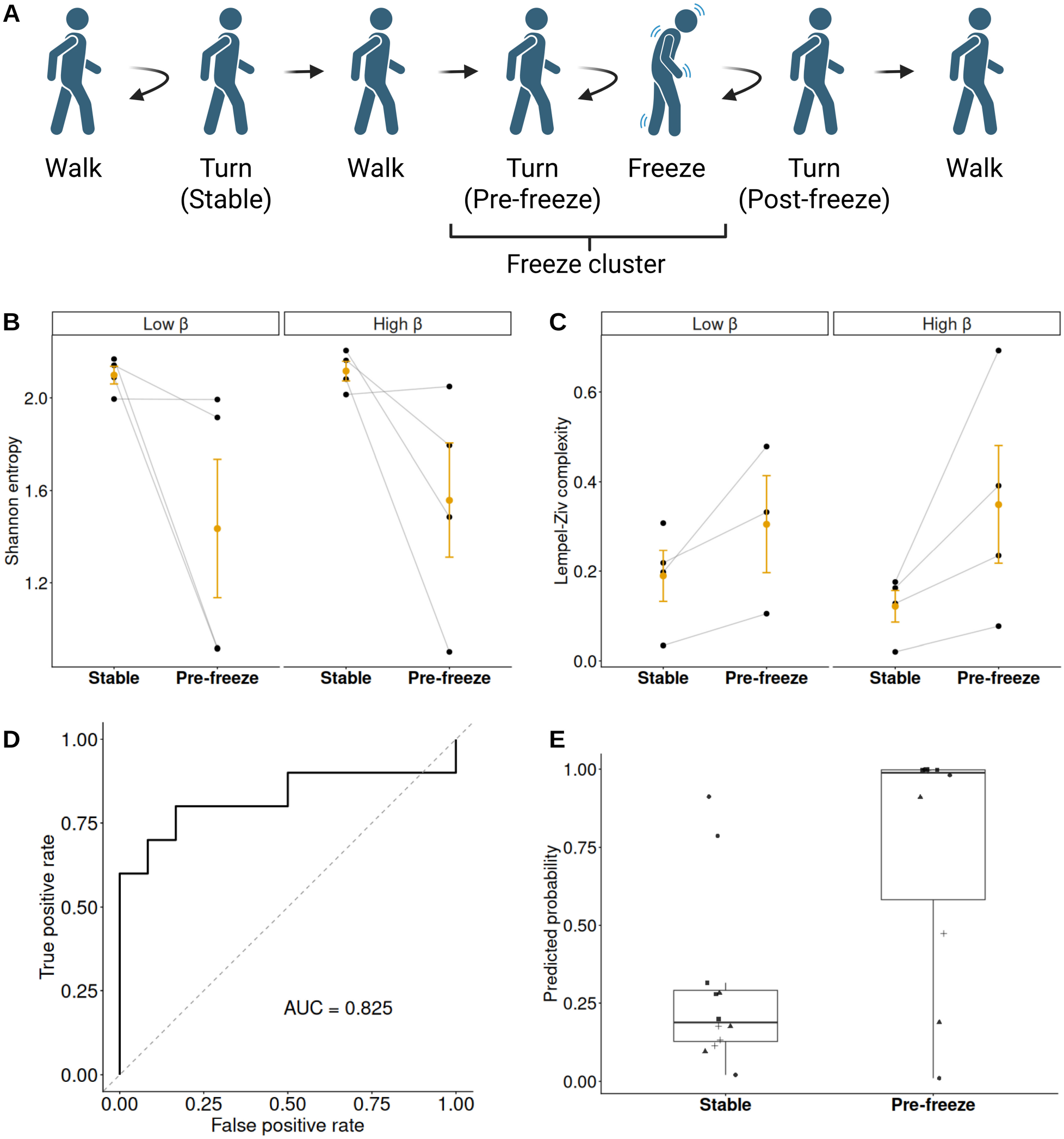
Pre-freeze dynamics and prediction of freezing onset (A) Schematic of the behavioural structure used for analysis. Turns preceding freezing within turn-freeze clusters were defined as pre-freeze turns and compared with stable turns. (B) Shannon entropy of burst timing is reduced during pre-freeze turns compared to stable turns in both low and high β bands. Lines indicate within-patient values; orange markers show mean ± SEM. (C) Lempel-Ziv complexity is increased during pre-freeze turns, consistent with increased temporal structure. One patient lacked sufficient pre-freeze data in the low beta band and was excluded from that comparison. (D) ROC curve for leave-one-patient-out classification of pre-freeze vs stable turns using entropy. (E) Predicted probabilities from the entropy-based classifier demonstrate separation between stable and pre-freeze turns across patients.

To assess predictive performance, a leave-one-patient-out logistic regression classifier was trained to distinguish pre-freeze from stable turns. Using Shannon entropy alone, the model achieved strong classification performance (AUC = 0.825, accuracy = 0.773), reliably separating pre-freeze from stable turns across patients (Figure 1D). Predicted probabilities were higher for pre-freeze turns within each patient (Figure 1E). Band-specific models yielded similar performance (low beta: AUC = 0.815; high beta: AUC = 0.817). Combining low and high beta entropy did not improve performance (AUC ≈ 0.75), indicating that the predictive signal is shared across bands. A simple threshold-based classifier achieved comparable or higher performance (AUC = 0.858 for entropy), with thresholds derived within each training fold. Threshold classifiers based on band-specific entropy yielded similar results (AUC = 0.852 for low beta; 0.842 for high beta).

Alternative features showed weaker performance (Figure S1A). Models based on inter-burst variability (CV and MAD) achieved moderate-to-high performance (MAD AUC up to ≈ 0.82). Combining entropy with MAD did not improve AUC (entropy + MAD: AUC = 0.821 vs 0.825 for entropy alone), but increased accuracy (0.818 vs 0.773), showing improved threshold-level separation without better overall discriminability.

PAC showed modest and inconsistent differences across frequency bands. Alpha-low beta coupling was higher during pre-freeze turns compared to stable turns. Alpha-low beta coupling increased in pre-freeze vs stable turns (p ≈ 0.053), but this effect did not remain significant after correction for multiple comparisons. Consistent with this, PAC-based classifiers showed moderate performance (AUC ≈ 0.70-0.75) but did not outperform entropy. Combining entropy with PAC did not improve performance (entropy + PAC: AUC = 0.808).

Across models, entropy consistently provided the most robust and generalizable discrimination between pre-freeze and stable turns, and additional features did not yield meaningful improvements beyond this single measure.

### Freezing is associated with reduced variability and a rigid timing regime

Burst dynamics during freezing were compared with turning within behavioral episodes (cluster-level comparisons) to isolate state-dependent effects (Figure 2A-C). Inter-burst intervals became more regular during freezing, reflected by a reduction in the coefficient of variation (CV) across both bands (high beta: p = 3.7 × 10^-6^; low beta: p = 0.0016) and a reduction in MAD in high beta (p= 0.023; low beta: p = 0.47). These effects were consistent at the episode level, with greater variability across patients due to the limited sample size. In contrast, Shannon entropy did not differ between freezing and turning (high beta: p = 0.56; low beta: p = 0.09), indicating that entropy primarily reflects pre-freeze changes. Complementary measures revealed that this reduced variability is accompanied by increased structure: permutation entropy was modestly reduced (high beta: p = 0.030; low beta: p = 0.042), and Lempel-Ziv complexity increased in high beta (p = 0.0047; low beta trend: p = 0.057), consistent with increased temporal structure, while sample entropy showed no significant effect.

**Figure 2.**
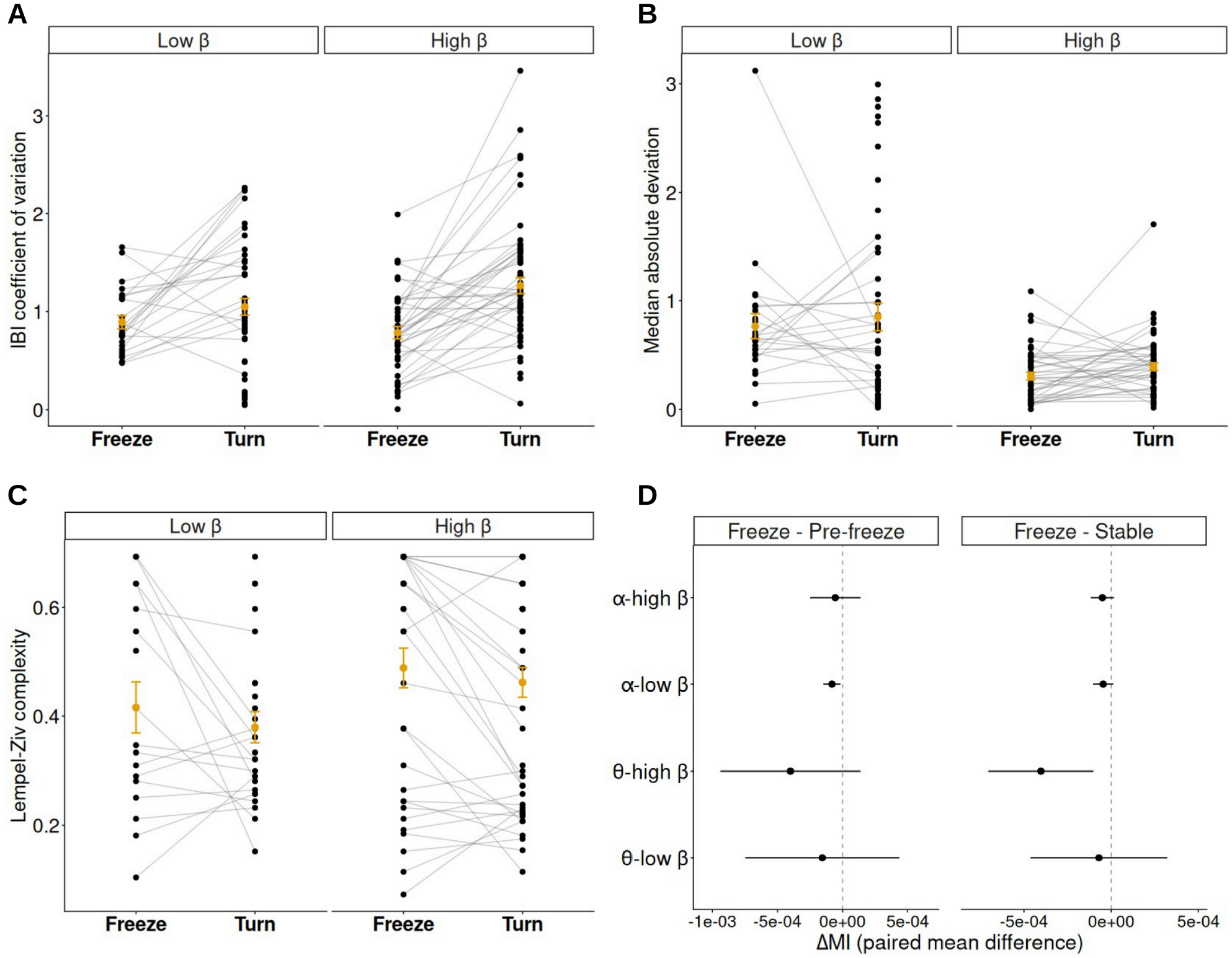
State-dependent changes during freezing Inter-burst interval CV (A) and MAD (B) are reduced and Lempel-Ziv complexity (C) is increased during freezing compared to pre-freeze turning at the cluster level. (D) PAC differences (ΔMI) for freezing relative to pre-freeze turns and stable turns across frequency bands. Points show mean differences ± SEM.

To assess whether these features capture the frozen state, a leave-one-patient-out classifier was trained to distinguish freezing from pre-freeze turns within the same episodes. Variability-based measures provided the strongest discrimination, with inter-burst interval CV and MAD yielding the highest performance (CV: AUC = 0.626; MAD: AUC = 0.617; combined CV + MAD: AUC = 0.629). In contrast to the pre-freeze comparison, Shannon entropy showed weaker performance (AUC = 0.596), demonstrating limited sensitivity to the frozen state itself. Measures of temporal structure, including permutation entropy (AUC = 0.585) and Lempel-Ziv complexity (AUC = 0.592), provided modest discrimination but did not exceed variability-based features.

PAC showed frequency-specific differences across state comparisons (Figure 2D). Alpha-low beta coupling was reduced during freezing relative to pre-freeze turns, while theta-high beta coupling was reduced during freezing relative to stable turns (p = 0.0528 for both). Decreased theta-high beta coupling was observed between freezing and stable turns (p = 0.032). However, these effects did not remain significant after correction for multiple comparisons, and no consistent differences were observed between freezing and pre-freeze turns. PAC-based classifiers showed high apparent performance (AUC ≈ 0.859), particularly for alpha-low beta coupling, though these results should be interpreted cautiously given the smaller sample size (Figure S1B). Combining Lempel-Ziv complexity with PAC (alpha-low beta) improved classification performance (AUC ≈ 0.86), suggesting that temporal structure and cross-frequency coordination capture complementary aspects of the freezing state.

These findings indicate that freezing is characterized by reduced inter-burst variability, accompanied by increased temporal structure and decreased coordinated cross-frequency activity, distinct from entropy-based changes that precede freezing.

## Discussion

This study identifies reduced global variability of STN beta burst timing as a robust feature preceding FOG. Shannon entropy distinguished pre-freeze from stable turning and provided strong predictive performance. Reduced Shannon entropy reflects more constrained and predictable burst timing, suggesting that STN activity explores a narrower range of temporal patterns before freezing.

This suggests a reduction in the accessible temporal state space of STN activity. This complements prior work linking prolonged beta bursts to freezing and showing improvement with adaptive deep brain stimulation(8,17). Previous work using this dataset demonstrated that low beta power increases during freezing and supports classification of locomotor states with moderate accuracy (∼50-75%)(16). However, these amplitude-based features did not distinguish between pre-freeze, freeze, and stable states. Notably, burst timing entropy provided robust discrimination of pre-freeze states. These results suggest pathological bursting emerges from a constrained temporal repertoire.

During freezing, burst timing became more regular, reflected by reductions in CV and MAD and accompanied by increased structure in coarse-grained representations (Lempel-Ziv complexity). In contrast to the pre-freeze condition, entropy did not reliably distinguish freezing from pre-freeze turns, indicating that it primarily captures changes preceding freezing.

PAC showed heterogeneous, frequency-specific and did not outperform burst-based features, indicating that PAC reflects secondary rather than primary network changes underlying freezing. Notably, reduction in PAC during freezing suggests diminished cross-frequency coordination within STN activity, consistent with a breakdown of network integration. This aligns with prior work showing that freezing is associated with disrupted cortical-subcortical communication(4) and abnormal cortical PAC patterns(15). These findings suggest that cross-frequency coupling is dynamically reconfigured across the transition into freezing rather than increasing monotonically.

These data suggest a two-stage model: a reduction in global temporal variability precedes freezing, followed by a shift to a more regular and structured regime during freezing. Within this framework, the pre-freeze reduction in entropy may reflect a shift toward conditions that favour the emergence of prolonged or pathological beta bursts. This interpretation suggests that freezing may arise when temporally flexible encoding of gait-related activity in the STN becomes constrained, with entropy providing a simple and robust predictor of its onset and a potential biomarker for identifying pre-freeze neural states.

## Acknowledgements

The research was supported by the National Institute for Health Research (NIHR) Oxford Biomedical Research Centre (BRC). The views expressed are those of the author(s) and not necessarily those of the NHS, the NIHR or the Department of Health.

## Data availability

All MATLAB and R analysis scripts relevant to this study are available at https://github.com/cabeaudoin/fog_entropy. Original data can be found at Thenasie *et al*. (https://zenodo.org/records/6867624)(16).

