## Supplementary Material for "Reduced entropy of subthalamic beta bursts predicts freezing of gait in Parkinson’s disease"

### Methods

*Dataset and recording structure*

We analysed subthalamic nucleus (STN) local field potential recordings during gait from the Thenaisie et al. dataset(1), restricted to the four participants included in the current study. For each participant and session, the MATLAB preprocessing pipeline loaded the session-specific LFP file and the corresponding behavioural annotation file, extracted the numeric LFP matrix from the source structure, oriented the matrix into a consistent time-by-channel format when needed, and matched the behavioural state vector to the LFP sampling grid if their lengths differed. Sampling frequency was 250 Hz. The downstream analyses were therefore performed on a session-aligned LFP time series with synchronised behavioural annotations.

*LFP preprocessing and channel selection*

For each session, the analysis channel was selected from the available LFP channels as the channel with the highest beta-power proxy. The proxy was computed as the mean squared amplitude of the 13-30 Hz band-pass filtered signal, after orienting the raw matrix appropriately and handling both vector and matrix inputs. The selected signal was demeaned before further processing. Band-pass filtering was implemented using a fourth-order Butterworth filter with zero-phase forward-reverse filtering. This same channel-selection strategy was used for the burst analysis and the PAC analysis.

*Burst detection*

Burst detection was performed separately in low beta (13-20 Hz) and high beta (20-30 Hz). For each band, the selected LFP was filtered, the analytic signal was obtained using the Hilbert transform, and burst amplitude was defined from the magnitude of the Hilbert envelope. Bursts were identified as contiguous periods during which the envelope exceeded the 75th percentile of the band-specific envelope distribution within each session. To avoid brief threshold crossings, only bursts lasting at least 2/f_low were retained, corresponding to approximately 154 ms for low beta and 100 ms for high beta. This approach is consistent with commonly used definitions of beta bursts in Parkinson’s disease LFP recordings(2,3). For each retained burst, we exported the start and end sample indices, start and end times, midpoint time, duration, peak amplitude, mean amplitude, integrated envelope, and behavioural state at both onset and midpoint.

*Behavioural state annotation and episode definitions*

Continuous state labels were converted into bouts of constant state by identifying changes in the state code. Each bout was assigned a unique bout identifier. For the turn-freezing analyses, consecutive TURN and FREEZE bouts were grouped into turn-freeze clusters. Within each cluster, TURN bouts preceding the first FREEZE bout were labelled pre-freeze turn, whereas TURN bouts not adjacent to FREEZE on either side were labelled stable turn. FREEZE bouts were labelled freeze. Unknown states were excluded. For the state analysis, TURN and FREEZE bouts were compared within the same cluster, preserving the turn-freeze sequence structure that defines the behavioural transition. Where needed, bursts were linked to state bouts via the burst midpoint.

*Burst-timing and complexity features*

For each patient, session, band, and behavioural condition, burst count, burst rate, total time in burst, mean peak amplitude, and mean burst duration were calculated. Burst timing was summarised using burst midpoint times, from which inter-burst intervals (IBIs) were derived as consecutive midpoint differences. From the IBI sequence, we computed the mean, standard deviation, median absolute deviation (MAD), coefficient of variation (CV), lag-1 autocorrelation, sample entropy, permutation entropy, and Lempel-Ziv complexity. Shannon entropy(4) was computed from the distribution of burst midpoint times using five equal-width bins spanning the observed time range, which was selected to balance robustness to noise and limited data with simplicity and interpretability for potential real-time clinical use. Sample entropy(5) was computed with embedding dimension $m = 2$ and tolerance $r = 0.2 \times SD$ of the sequence; permutation entropy(6) used embedding dimension $m = 3$ and delay = 1; and Lempel-Ziv complexity(7) was calculated after binarizing the IBI sequence relative to its median. These features were computed only when the sequence length was sufficient for the relevant statistic.

*State-comparison analysis*

The primary state comparison used cluster-level paired data, because the behavioural unit of interest was the TURN/FREEZE cluster rather than individual bursts. For each cluster, TURN and FREEZE values were paired within the same patient, session, and cluster identifier. Paired comparisons were performed using Wilcoxon signed-rank tests. For visualisation, paired cluster-level values were plotted with within-cluster lines connecting TURN and FREEZE. Patient-level summaries were additionally computed by averaging across clusters within each patient and band, but these were used only for secondary summaries and sanity checks rather than primary inference. This distinction was important because the inferential unit for the main state comparison was the cluster, whereas the display unit in some plots was the patient.

*PAC extraction and quantification*

PAC was computed in a separate MATLAB pipeline using the same session files and the same channel-selection rule. After channel selection and demeaned preprocessing, the state vector was resampled to match the LFP length when necessary using nearest-neighbour interpolation, preserving discrete labels. PAC was estimated only within bouts labelled pre-freeze turn, stable turn, and freeze. The phase band was 8-12 Hz and the amplitude band was 20-30 Hz. Phase-amplitude coupling was quantified with Tort's modulation index(8): the Hilbert phase of the phase-band signal was binned into 18 bins over $[-\pi,\pi]$, mean amplitude was computed within each bin, and the normalized divergence of this amplitude distribution from uniformity was used as the PAC index. PAC was computed in 2.0 s windows stepped every 250 ms, and the resulting tables included patient, session, episode type, bout identifier, window start and end indices, center time, phase band, amplitude band, and modulation index.

*PAC statistical analysis*

PAC contrasts were summarised at the patient/session/episode level by averaging MI within each episode type for each PAC file. Three contrasts were examined: pre-freeze turn versus stable turn, freeze versus pre-freeze turn, and freeze versus stable turn. Paired Wilcoxon signed-rank tests were used for the primary comparisons, and p-values were adjusted across the four PAC band combinations within each contrast using the Benjamini-Hochberg procedure. This provided a conservative summary of the frequency-specific PAC effects while preserving the matched structure of the episode-level data.

*Predictive modelling*

Predictive analyses used a leave-one-patient-out logistic regression framework. For each fold, one patient was held out as the test set and the remaining patients were used for training. Predictor variables were z-scored using the training-set mean and standard deviation, then applied to the held-out patient using the same scaling parameters. Models were fit with logistic regression and evaluated on the held-out patient using predicted probabilities. Performance was summarized with ROC AUC, accuracy, and balanced accuracy. In addition to multivariable models, we also fit single-feature threshold classifiers. Thresholds were selected on the training set using Youden's index from the ROC curve and then applied to the held-out patient. This framework was used for both pre-freeze versus stable-turn decoding and freeze versus pre-freeze-turn decoding. The model grid included entropy-only, band-specific entropy, entropy combined with PAC, entropy combined with variability features, PAC-only, and full multivariable models.

*Statistical software and implementation*

Signal preprocessing, burst detection, bout assignment, and PAC export were performed in MATLAB (MathWorks; version R2025B). Downstream aggregation, statistical testing, plotting, and decoding were performed in R using dplyr, tidyr, purrr, ggplot2, pROC, lme4, and lmerTest(9). Missing values were handled by complete-case analysis within each comparison or model. For the mixed-effects analyses used in the pre-freeze versus stable-turn comparison, turn role and beta band were included as fixed effects and patient as a random intercept. Bout duration was additionally included as a covariate in a sensitivity analysis.

| Numbering in original manuscript | Numbering in code | Medication status | Age | Sex | Disease duration (years) | Dominant symptoms | Score ON/OFF |
|---|---|---|---|---|---|---|---|
| E2 | PD06 | OFF (>12 h withdrawal) | 55 | M | 7 | AR | 30 / 7 (UPDRS III) |
| E3 | PD09 | Partial OFF (residual) | 67 | M | 11 | AR+T | 37 / 18 (MDS-UPDRS III) |
| P1 | PD19 | Partial OFF (residual) | 58 | F | 9 | AR+T | 17 / 6 (MDS-UPDRS III) |
| P4 | PD13 | ON medication | 60 | M | 7 | AR+T | 45 / 33 (ON 1 h / OFF 18 h, MDS-UPDRS III) |

Table S1. Clinical characteristics of participants included in the study.
Data reproduced from Thenaisie *et al.* (1). UPDRS part III scores are reported as ON/OFF where available. Medication status reflects clinical state at the time of recording.

| | | # Bouts | | | Duration (s) | | |
|---|---|---|---|---|---|---|---|
| Numbering in original manuscript | Numbering in code | FREEZE | STABLE TURN | PRE-FREEZE TURN | FREEZE | STABLE TURN | PRE-FREEZE TURN |
| E2 | PD06 | 42 | 29 | 35 | 3.07 ± 1.31 | 8.58 ± 1.07 | 2.62 ± 2.03 |
| E3 | PD09 | 72 | 68 | 19 | 7.82 ± 7.85 | 35.77 ± 27.58 | 14.18 ± 7.37 |
| P1 | PD19 | 31 | 10 | 6 | 1.87 ± 1.62 | 21.36 ± 13.59 | 6.59 ± 4.25 |
| P4 | PD13 | 24 | 42 | 15 | 1.16 ± 0.77 | 4.43 ± 0.83 | 1.55 ± 0.90 |

Table S2. Summary of freezing-related behavioural bouts for each participant. Number and duration (mean ± SD, seconds) of freeze bouts, stable turns, and pre-freeze turns included in the analysis.

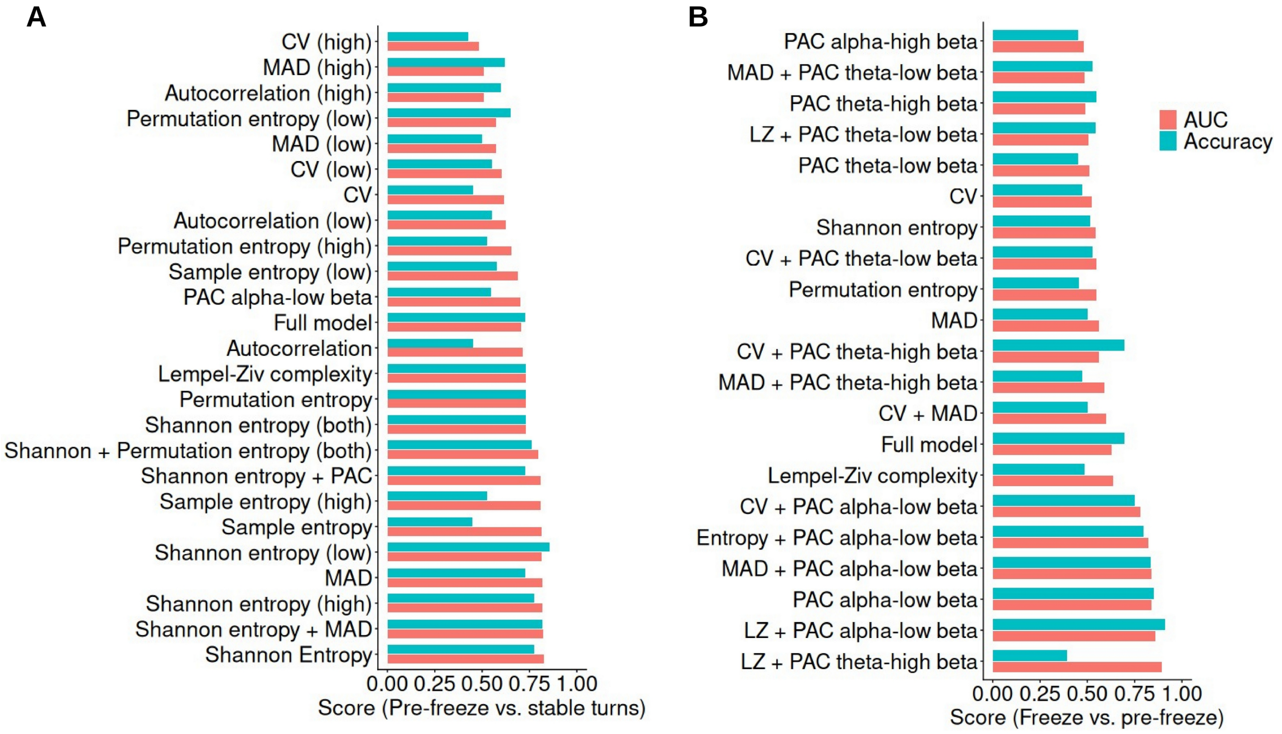


Figure S1. Model comparison for decoding performance
(A) Decoder performance (AUC and accuracy) for distinguishing pre-freeze vs stable turns across features and models. (B) Decoder performance for distinguishing freezing vs pre-freeze turns. Bars show AUC and accuracy for leave-one-patient-out classifiers.
